# Multiyear Improvement In Batting Skills Following Targeted Perceptual Cognitive Training In Softball

**DOI:** 10.1101/2024.08.29.610321

**Authors:** Jordan Muraskin, Jason Sherwin

## Abstract

**Background:** Making quick, accurate decisions is crucial in competitive sports like softball, where perceptual-cognitive skills can significantly impact on-field performance. This study evaluates the long-term effectiveness of a targeted perceptual-cognitive training program, delivered through the uHIT platform, on improving batting performance in collegiate softball players.

**Methods:** A longitudinal analysis was conducted on data collected from both NCAA and NAIA softball teams over multiple seasons. The study used Bayesian statistical methods to assess the impact of cognitive training on on-base plus slugging percentage (OPS). The analysis incorporated weighted models to account for variability in games played and differences between teams, and the influence of team and year-division effects was considered. Key metrics, including Decision AUC and Response Time, were tracked to evaluate perceptual-cognitive improvements.

**Results:** The results demonstrated significant improvements in OPS for teams that participated in the cognitive training intervention, with the weighted models indicating a substantial effect of the training on performance. Notably, the intervention was most effective in teams with higher training intensity, as evidenced by the permutation test results. The Bayesian analysis also revealed that the intervention led to statistically significant improvements in decision-making and response times, translating into enhanced on-field performance.

**Conclusion:** The findings support the effectiveness of perceptual-cognitive training in improving real-world athletic performance in softball. The uHIT platform, as an ecologically valid training tool, has demonstrated its potential to serve as a critical component of athletic development programs. Future research should explore the long-term retention of these cognitive gains and their application across different sports and competitive levels.

## Introduction

In competitive sports, particularly baseball and softball, making quick, accurate decisions is crucial for success. As the game unfolds, athletes must rapidly interpret visual information and translate it into action, whether recognizing the type of pitch thrown or deciding when to swing. Over the past few decades, there has been a growing interest in the role of cognitive training in enhancing these perceptual and decision-making skills (Mann et al., 2007). However, while cognitive training programs have shown promise in laboratory settings, there is still a need to understand how these interventions translate into real-world athletic performance, particularly over extended periods (Fransen, 2024).

Understanding how hitters see and make decisions led to the creation of the uHIT platform, a computerized training tool that simulates real-life pitch recognition scenarios (Muraskin et al., 2016; Muraskin et al., 2017; Muraskin et al., 2015; J. Sherwin et al., 2012, J. S. Sherwin et al., 2015). Developed by deCervo, the uHIT system offers an ecologically valid environment where athletes can hone their perceptual decision-making skills under conditions that closely mirror those experienced in actual gameplay. This platform has been used at professional and collegiate levels to improve cognitive skills such as pitch recognition and decision-making speed, which are critical for success in baseball and softball.

This study aims to show that targeted cognitive training can have the long-term perceptual-cognitive effects on collegiate softball players. Unlike previous studies that primarily focused on short-term outcomes, this research takes a longitudinal approach, assessing the impact of cognitive training over multiple seasons. By analyzing data from both NCAA and NAIA teams, we seek to determine whether improvements in cognitive skills, as measured by metrics such as Decision AUC and Response Time, lead to sustained enhancements in on-field performance, specifically in terms of on-base plus slugging percentage (OPS).

Cognitive training, which involves exercises designed to improve mental processes like attention, memory, and decision-making, has been shown to enhance perceptual-cognitive expertise in athletes (Farrow & Abernethy, 2002; Fadde, 2016). These skills are particularly relevant in baseball and softball, where the ability to quickly and accurately perceive and react to a fast-moving ball can distinguish elite players from their peers. The theory of cognitive transfer suggests that skills developed in one context can improve performance in another, provided sufficient overlap exists in the cognitive processes involved (Barnett & Ceci, 2002). However, the extent to which this transfer occurs in real-world sports settings remains inconclusive (Fransen, 2024).

This study employs a Bayesian statistical framework to analyze the data and build upon the existing literature. Using Bayesian methods allows for a more nuanced interpretation of the data, particularly in accounting for game variability and differences between teams. Additionally, the application of weighted models in this study addresses a standard limitation in sports research, where unequal participation rates can introduce bias. By incorporating these advanced statistical techniques, we aim to provide a more robust analysis of the effectiveness of cognitive training in enhancing softball performance.

This study’s findings can inform the design of more effective training programs in baseball, softball, and other sports in which cognitive skills play a critical role. If cognitive training can have a lasting impact on performance, it could become an essential component of athletic development programs at all levels of competition. This research thus contributes to a growing body of evidence supporting integrating cognitive training in sports.

## Methods and Data

### Perceptual Training with uHIT

uHIT data is made by playing one of two uHIT modules. Pitch Recognition is the first. In this modified go/no-go paradigm, the player is shown a visual cue before seeing a simulated pitch trajectory. If the class of that cue matches the trajectory class, then the player must ‘go’ as quickly as possible. The response is deemed incorrect if the player goes after the trajectory completes. The player must withhold a physical response (‘no-go’) if there is no match. For instance, if the cue is from the ‘fastball’ class and the trajectory is also from the ‘fastball’ class, the player must ‘go’ as quickly as possible and before the trajectory completes. With a different cue, the player must withhold a response, for instance if the trajectory had been of the ‘curveball’ class. For softball, the pitch types in the Pitch Recognition module are Riseball, Curveball, and Dropball.

The second uHIT Game is Zone Recognition. This is also a modified go/no-go paradigm. Here, though, there is no pre-trajectory cue. Instead, the player is shown an outline of the strike zone before the pitch trajectory comes. The player must ‘go’ as quickly as possible if he expects the trajectory to terminate in the strike zone. He is to withhold a response if he expects it not to. The pitches in the Zone Recognition module are Riseball, Curveball, Dropball, and ChangeUp.

Together, these two tasks assess the crucial abilities of baseball and softball hitters. The first task, Pitch Recognition, assesses a player’s ability to predict and identify pitch types. The second task, Zone Recognition, assesses a player’s ability to predict whether a pitch will be a strike.

We developed an intervention program for pitch identification in softball hitting. We based the program on an initial assessment (“Assessment”) of pitch recognition ability. As discussed in prior reports (Muraskin et al., 2016; Muraskin et al., 2017; Muraskin et al., 2015; J. Sherwin et al., 2012, J. S. Sherwin et al., 2015), pitch recognition is a perceptual decision-making skill that is a time-pressured go / no-go decision. The Assessment measures performance at this skill with such behavioral measures as area under the receiver-operator curve (“AUC”), reaction time, and other metrics. We found that completion of the Assessment in an elite training environment is highly dependent on the number of trials and the task’s difficulty. From prior data, we determined that n = 30 trials repeated in two (2) blocks balances attentional demands on the participant with the sample size needed to generate an intervention program. Also, for elite players, we determined that a mean pitch speed of 48 miles per hour in softball allows enough of a challenge while not making the Assessment too difficult. If the Assessment is too lengthy or hard, we risk greatly diminishing participation from elite players.

Our intervention program for pitch recognition (also called “uHIT Custom”) modifies two variables that balance 1) the difficulty of the task and 2) the length of the training. We modify these three variables together based on an initial Assessment and subsequent re-assessments (“Re-Assessment”) to maintain remote participation and continued progress in the difficulty-changing program.

### Subjects

Three collegiate softball teams were recruited to participate in the study. One of these (NAIA Team #1) is a Division I team from the National Association of Intercollegiate Athletics. The other two are Division I (NCAA Team #1 and NCAA Team #2) from the National Collegiate Athletic Association. NAIA Team #1 was recruited in the Fall 2020 and used the uHIT Platform starting September 2020. NAIA Team #1 used uHIT during the fall and spring seasons through 2024. Throughout the study, 52 hitters completed some training on the uHIT platform. NCAA Team #1 was recruited in the Fall of 2021. The team continued to use uHIT for the fall and spring seasons through 2024. For the study, 28 players completed some training on the uHIT Platform. NCAA Team #2 was recruited in the Fall of 2020. The team used uHIT in the Fall of 2020 and Spring of 2021. In that season, 28 players completed some uHIT Training. The coaches on each team assigned uHIT to players as a practice tool.

### NCAA Data

Division I Team softball hitting data was collected from stats.ncaa.org for the years 2015 through 2024. Team stats included basic stats from each season. These included Games Played (GP), At-Bats (AB), Runs, Hits (H), Doubles (2B), Triples (3B), Total Bases (TB), Home Runs (HR), Intentional Walks (IBB), Walks (BB), Hit By Pitch (HBP), and a sacrifice fly (SF).

### NAIA Data

Softball hitting data for NAIA Division I teams were collected from two sources: dakstats.com and naiastats.prestosports.com. The NAIA changed its official stats provider from Dakstats to Presto Sports after the 2021 season. Data was collected from 2015 to 2021 from Dakstats on all teams and from 2022 to 2024 from Presto Sports.

### On-Base Plus Slugging (OPS) Calculation

The On-Base Plus Slugging (OPS) is a number that summarizes a hitter’s offensive output from available hitting data. It’s meant to combine how well a hitter can reach base with how well he can hit for average and power (www.mlb.com/glossary). We manually calculated the OPS for each team in each season by first calculating the On-base Percentage (OBP) and Slugging Percentage.

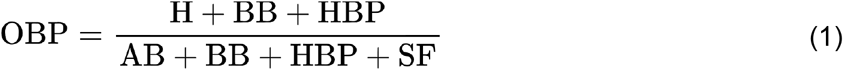

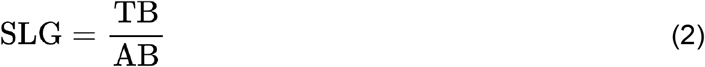

where **TB** is calculated as:**TB=(H-2B-3B-HR)+2.2B+3.3B+4.HR**

OPS is then calculated as:

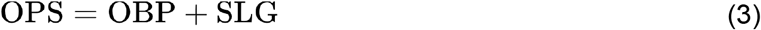

### Data Preprocessing

The dataset comprised collegiate softball team statistics spanning multiple years, focusing on each team’s OPS metric. Initially, the data were sorted chronologically by team and year to ensure proper sequencing of observations. For each team, the OPS of the previous year (**OPS**_prew_) was calculated using a lag function within each division, allowing us to track performance changes over time.

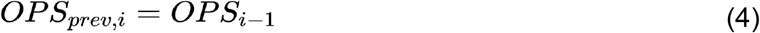

The difference between a team’s previous year’s OPS and the previous year’s average OPS was then calculated and standardized. This was done to account for variations in performance across different years and divisions. The change in OPS (Δ**OPS**) was calculated as the difference between the current year’s OPS and the previous year’s OPS and then standardized to ensure numerical stability. In addition, **OPS**_prew_ was also standardized by subtracting the mean of OPS and dividing it by the standard deviation of OPS.

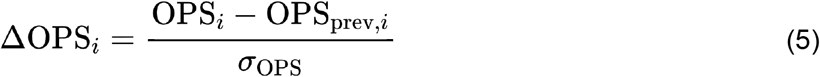

The Year and Division variables were encoded as categorical variables to facilitate the inclusion of year-specific and division-specific effects in the model. Intervention periods were defined for specific teams and encoded as a binary variable, with 1 indicating the application of an intervention and 0 otherwise.

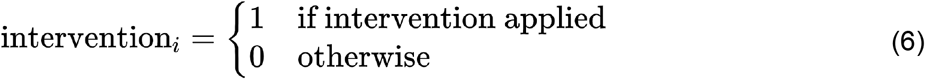

Weights were calculated based on the number of games played and normalized to ensure that observations with more games played had a greater influence on the model.

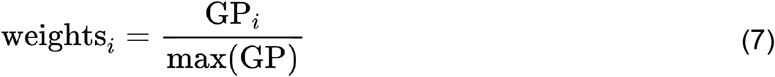

### Model Specification

We constructed four different models to assess the impact of the intervention on team performance and compare their predictive performance:

#### 1. Weighted Model with Intervention

This model included the intervention effect and weighted the observations based on the number of games played. Weighting the observations by the number of games played helps to account for variability in the sample size, giving more influence to observations with more games. The linear predictor for this model is:

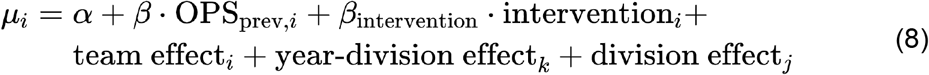

*α* is the model intercept and is modeled as a normal Gaussian:

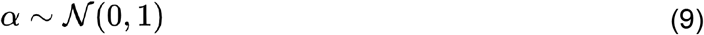

*β* is the coefficient applied to the previous year’s OPS for the team and is also modeled as a normal distribution:

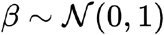

*β*_intervention_ is the coefficient applied to the intervention variable. It is also modeled as a normal distribution. *β*_intervention_will capture if the cognitive training affects the change in OPS.

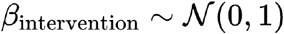

#### 2. Weighted Null Model

This model excluded the intervention effect but still weighted the observations by the number of games played. This model serves as a baseline to assess the added value of the intervention effect in the weighted model. The linear predictor for this model is:

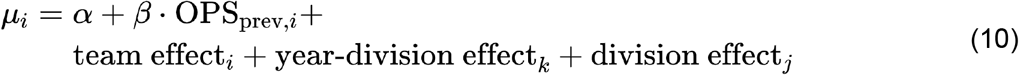

#### 3. Non-Weighted Model with Intervention

This model included the intervention effect but did not add weights to the observations. It allows us to compare the impact of weighting on the model’s performance and understand the effect of the intervention without considering the number of games played. The linear predictor for this model is the same as the Weighted model.

#### 4. Non-Weighted Null Model

This model excluded the intervention effect and did not weight the observations. This model serves as a baseline to assess the added value of the intervention effect in the non-weighted model. The linear predictor for this model is the same as the Weighted Null model.

### Priors and Non-Centered Parameterization

Normal priors were specified for the global mean and standard deviation of team effects to account for the inherent variability among teams. The priors were defined as:

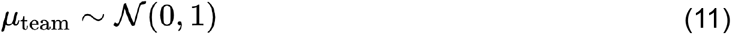

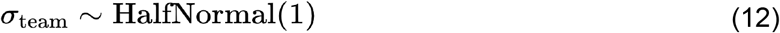

Team-specific effects were modeled using non-centered parameterization to improve sampling efficiency. The team effect for each team was modeled as follows:

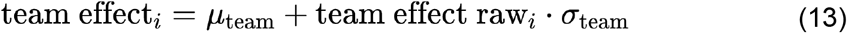

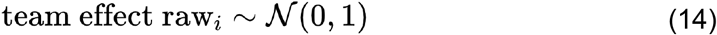

The team effect captures each team’s inherent performance level, accounting for differences not explained by other predictors. This allows the model to account for the fact that some teams may consistently perform better or worse than others due to factors not explicitly included in the model. Possible factors include coaching differences, financial resources, and recruiting ability, just to name a few.

Year-specific and division-specific effects were similarly modeled using non-centered parameterization to account for the variability across different years and divisions:

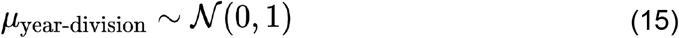

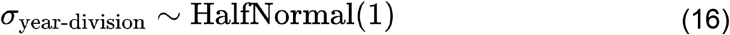

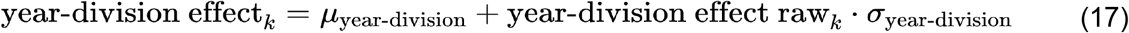

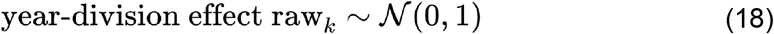

The year-division effect captures systematic differences in performance attributable to specific years and divisions, such as changes in league policies, environmental conditions, or competitive balance. Including these effects helps to control for external factors that influence team performance in a given year and division.

Division-specific effects were included to capture variability across divisions:

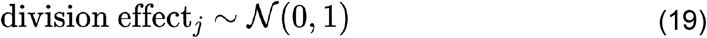

The likelihood of the observed change in OPS was modeled as follows:

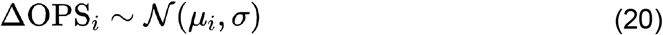

### Incorporating Weights in the Bayesian Model

To mathematically include the weights in the Bayesian model, the weights adjust the log-likelihood contributions for each observation. Specifically, the log-likelihood for each observation is multiplied by the corresponding weight, which reflects the number of games played for that season. This adjustment ensures that observations with more games played have a greater influence on the model:

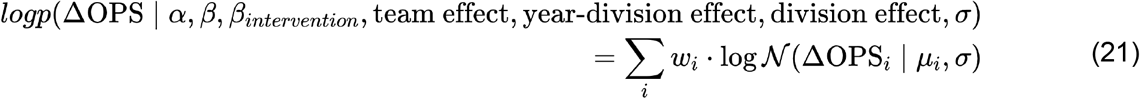

where 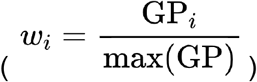 is the normalized weight for the i-th observation.

### Inference

Bayesian inference was performed using the No-U-Turn Sampler (NUTS) in PyMC (Abril-Pla et al., 2023) using Numpyro (Phan et al., 2019). Each model was run with 500 tuning steps and 1500 samples in 4 independent chains. The multiple chains were used to assess model convergence and stability.

### Model Comparison and Validation

To assess their predictive performance, the models were compared using Leave-One-Out (LOO) (Vehtari et al., 2016) cross-validation. The expected log predictive density (elpd_loo) was calculated for each model, and the models were ranked based on their elpd_loo values. Additionally, a permutation test was conducted to evaluate the significance of the intervention effect. The intervention labels were randomly shuffled, and the effect was recalculated 500 times to generate a distribution of permuted intervention effects. The p-value was determined as the proportion of permuted values greater than or equal to the observed effect. Each intervention reshuffle maintained a similar structure to the actual data. Each reshuffle had one team with a single intervention season, one with three consecutive seasons, and one with four consecutive seasons.

## Results

### uHIT Assessment and Training Results

Overall, NCAA Team #1 and NAIA Team #1 completed more training than NCAA Team #2. Figure 1 NCAA Team #1 (A) and NCAA Team #2 (B) display a relatively lower number of training pitches, with most batters receiving fewer than 2000 pitches. In contrast, NAIA Team #1 (C) exhibits a broader distribution, with some batters receiving as many as 20,000 training pitches. This figure highlights the variation in training intensity among the teams. Differences in coaching preferences and the number of seasons of use caused considerable differences between training regimens. NCAA Team #2 only used uHIT for a single season, while NAIA Team #1 trained with uHIT for four complete seasons (Fall and Spring) and NCAA Team #1 for three complete seasons.

**Figure 1:**
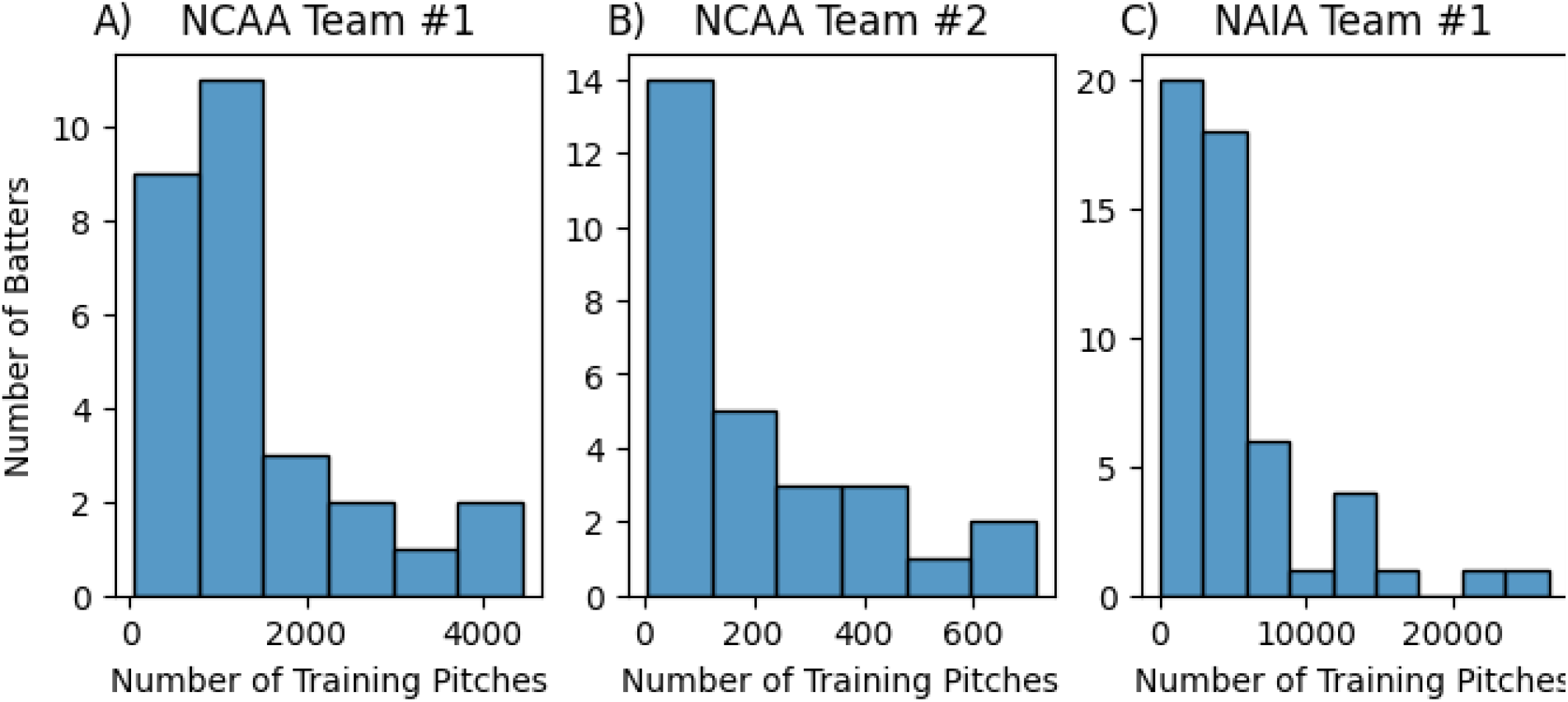
Histograms depict the number of batters and their corresponding number of training pitches for three different collegiate softball teams: (A) NCAA Team #1, (B) NCAA Team #2, and (C) NAIA Team #1. The x-axis represents the number of training pitches each batter received, while the y-axis shows the number of batters within each pitch count range.

Figure 2 plots assessment and Training performance changes. For Pitch Recognition (Figure 2A), all three teams improved their accuracy from their Assessment to the end of their training. Table 1 shows the t-values for the paired t-test comparing changes from Assessment to the final 10% of training sessions. All teams significantly (p=0.00004 for NCAA Team #1, p=0.017 for NCAA Team #2, p=4.9×10^−14^ for NAIA Team #1) improved their decision AUC during training. Only NAIA Team #1 significantly decreased (p=6.5×10^−12^) their Response Time.

**Table 1:** uHIT Team training results. Paired t-test t-value results comparing Assessment to the final 10% of sessions for each team broken down by decision AUC and Response Time. NCAA Team #2 focused their training on Pitch Recognition, so training results for Zone Recognition are N/A. * indicates p<0.05, *** indicates p<0.001.

|  | Pitch Recognition |  | Zone Recognition |  |
| --- | --- | --- | --- | --- |
|  | *AUC* | *RT* | *AUC* | *RT* |
| **NCAA Team #1** | 5.86*** | -0.51 | 1.77* | -1.43 |
| **NCAA Team #2** | 2.64* | -0.74 | N/A | N/A |
| **NAIA Team #1** | 10.37*** | -8.87*** | -0.97 | -4.82*** |

**Figure 2:**
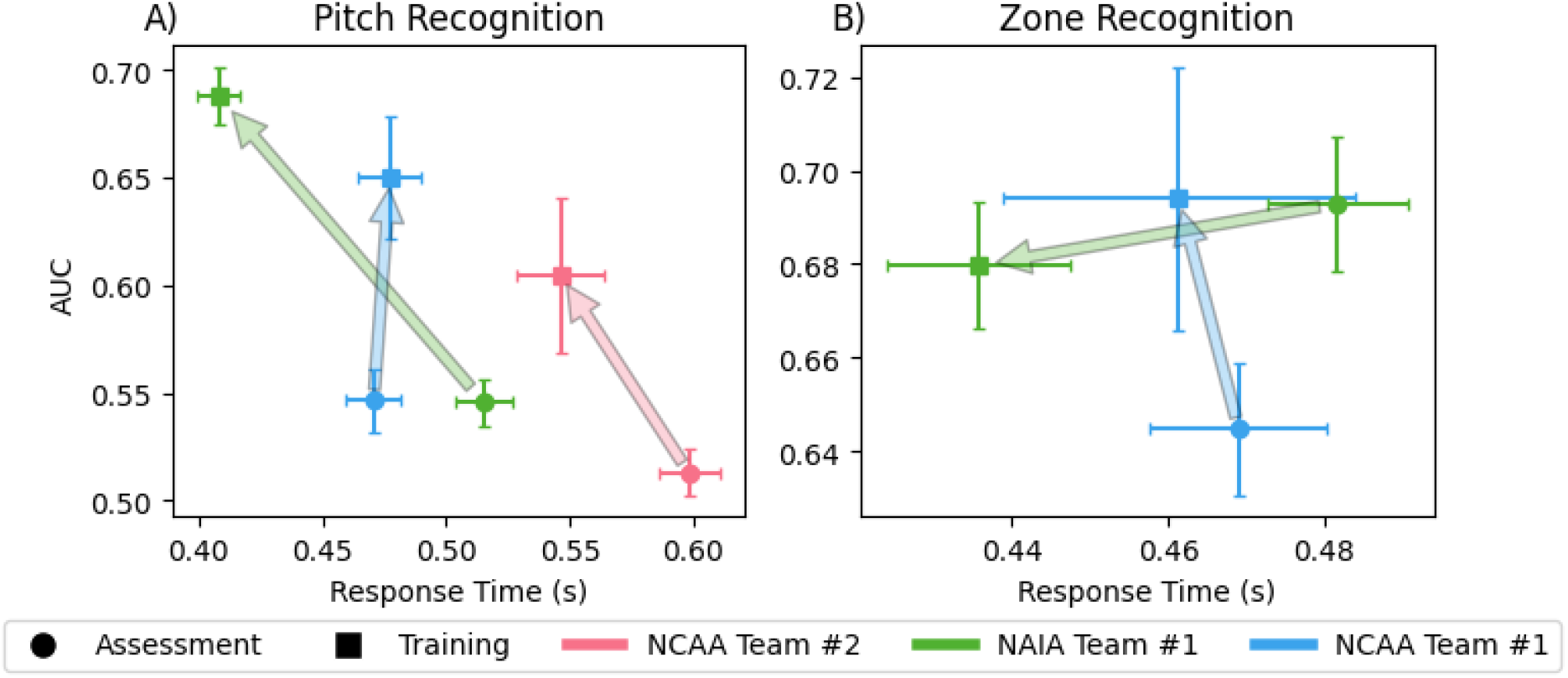
Relationship Between Response Time and AUC in Pitch and Zone Recognition for Three Collegiate Teams. (A) The graph shows the relationship between response time (in seconds) and Area Under the Curve (AUC) for Pitch Recognition across three collegiate teams: NCAA Team #1 (blue), NCAA Team #2 (red), and NAIA Team #1 (green). The arrows indicate the transition from assessment (circles) to training (squares). (B) The graph shows the relationship between response time (in seconds) and AUC for Zone Recognition across the same three teams. Error bars show the standard error in both the AUC and Response Time axes.

NCAA Team #2 focused exclusively on Pitch Recognition, so there wasn’t enough training on Zone Recognition to conduct a pre/post-analysis. While NAIA Team #1 did not increase their AUC during training, they significantly decreased their Response Time (p=0.00012). NCAA Team #1 significantly (p=0.048) improved their Zone Recognition decision AUC while slightly lowering their Response Times.

### On-Field NCAA and NAIA Data

On-field data for the NCAA and NAIA softball teams were compiled from 2015 through the 2024 seasons. There were 511 teams in the dataset with at least two seasons after removing three for incomplete data (stats were not available to compute OPS). Figure 3 shows the breakdown of NCAA and NAIA teams in the dataset by year, along with a histogram of the number of years each team is in the dataset. 426 of the teams were included in all ten years of the dataset. The calculation of team OPS is summarized in Figure 4A, which shows the yearly average of team OPS split for NCAA and NAIA. Figure 4B shows the season-to-season variability in the number of games teams play. The 2020 and 2021 seasons show a drop in games played due to the COVID-19 pandemic. The large in-season and season-to-season variability in the number of games played was a deciding factor in creating a model that weighted observations by the number of games played.

**Figure 3:**
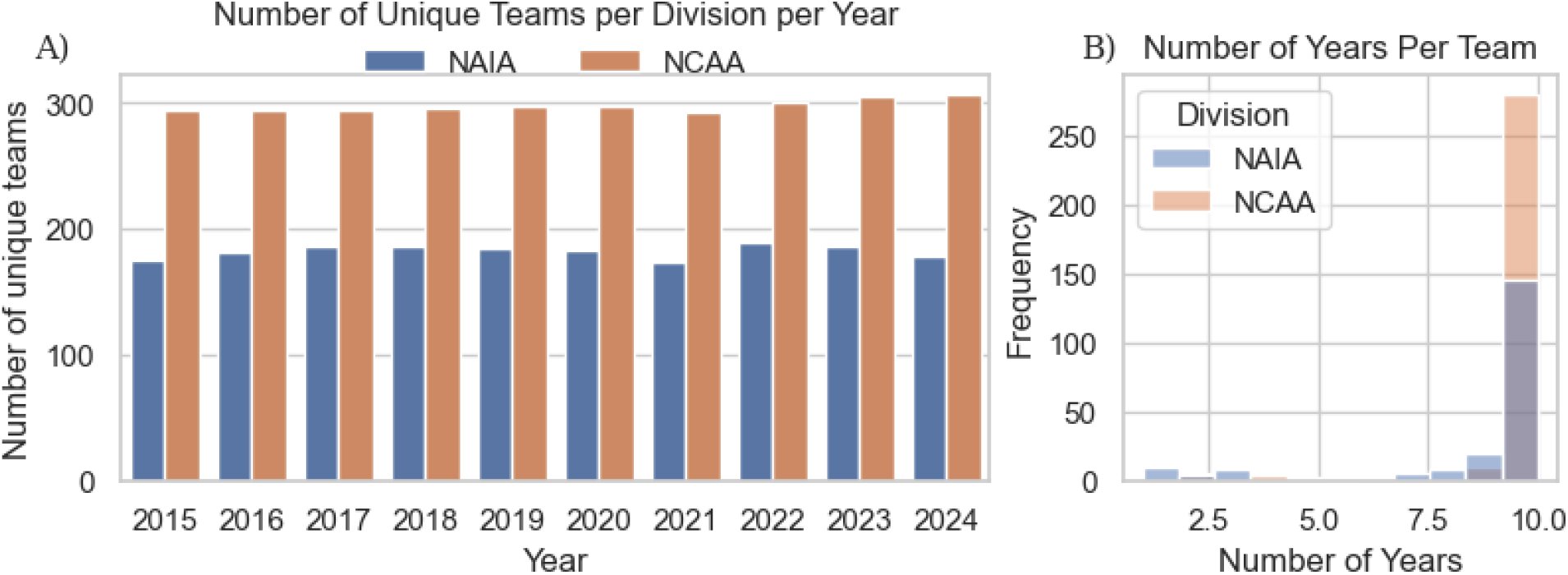
On-field dataset results showing the number of unique teams per season (A) and the distribution of the number of seasons for each team in the dataset (B).

**Figure 4:**
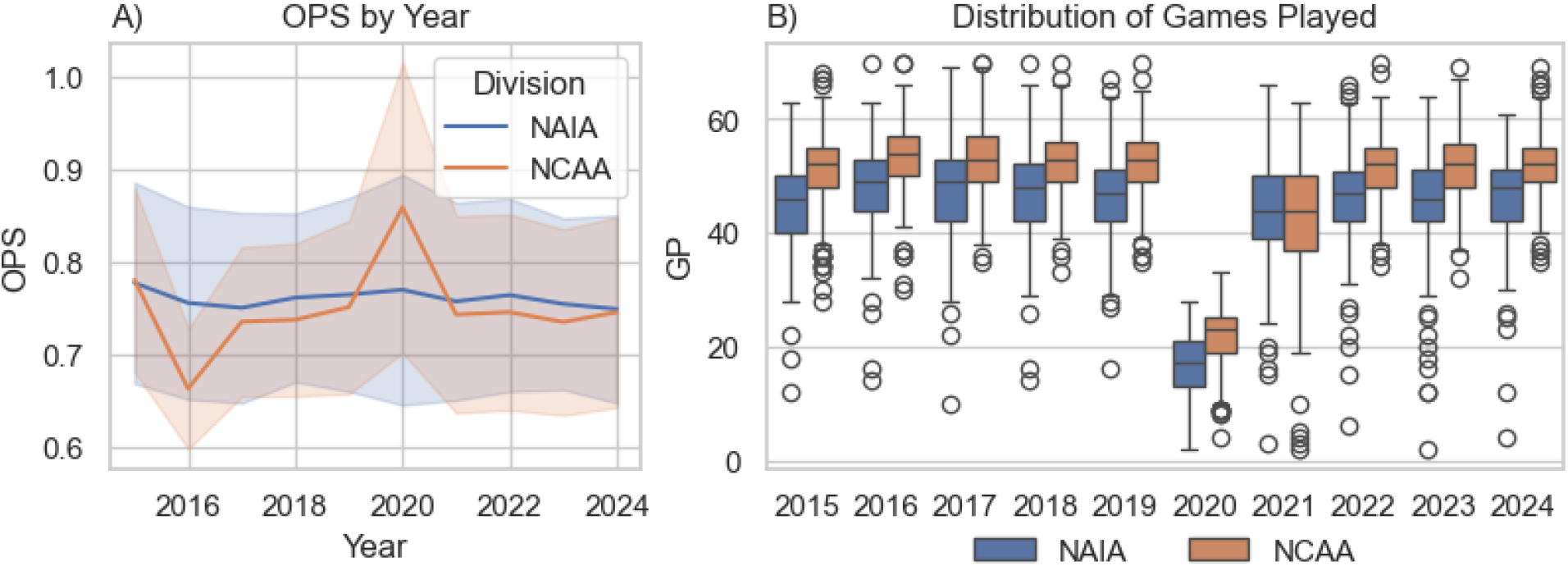
A) Season-to-season average OPS with standard deviation separated by NAIA and NCAA divisions. B) Shows the variability in games played by season: note the large drop in the 2020 season due to the Covid-19 pandemic.

The on-field OPS data for the three recruited teams is plotted in Figure 5. NAIA Team #1 data (Figure 5A) shows that the team’s OPS has been above the NAIA average for all seasons, with only two seasons below 0.900. The intervention years of 2021-2024 show that the team maintained their +0.900 OPS and did not regress to the mean. NCAA Team #1 OPS data (Figure 5B) shows more variability season-to-season. The intervention years show a linear trend of improvement. NCAA Team #2 OPS data shows similar variability to NCAA Team #1 but with only one intervention year. The best hitting year by OPS for NCAA team #2 was during the intervention year (2021 with an OPS of .931) and then returned to around the NCAA average.

**Figure 5:**
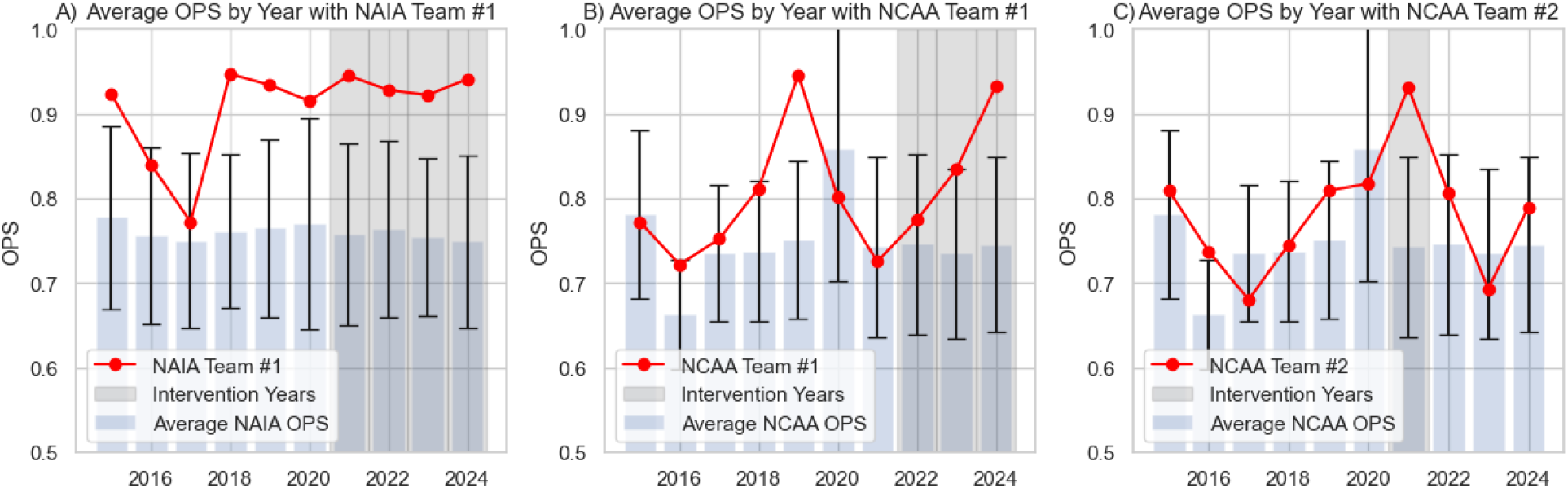
On-Field OPS season-to-season results for the 3 recruited teams. A) NAIA Team #1 OPS results (Red) compared to NAIA average OPS (blue bars) with black standard deviation error bars. B) NCAA Team #1 OPS season-to-season results with average NCAA OPS with standard deviation. C) NCAA Team #2 OPS season-to-season results. Intervention years are shaded in grey.

### Model Results

After preprocessing the above data, the four models were run with 500 tuning steps and 1500 samples across four independent chains. In the model, the team effect had 511 independent coefficients, and the year-division effect had 18. All models in our analysis successfully converged, as indicated by the Gelman-Rubin diagnostic (Gelman & Rubin, 1992), commonly referred to as 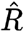. For all variables in each model, 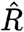 values were approximately equal to 1. The 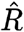 statistic is used in Bayesian modeling to assess the convergence of Markov Chain Monte Carlo (MCMC) simulations. Specifically, it compares the variance within each chain to the variance between chains. When 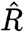 is close to 1, it suggests that the chains have converged to a standard distribution, indicating reliable posterior estimates.

To evaluate the predictive performance of our models, we conducted a Leave-One-Out Cross-Validation (LOO) analysis. The results of this analysis are summarized in Table 1. The model comparison revealed that the weighted model with the intervention effect had the highest expected log predictive density (elpd_loo) at -4532.24, making it the best-performing model among those tested. The weighted null model followed this model, with a slightly lower elpd_loo of -4535.44 and an elpd difference (elpd_diff) of 3.20. The small elpd_diff indicates that including the intervention effect in the weighted model provides a modest improvement in predictive performance. This modest improvement can be attributed to the small number of interventions compared to the rest of the dataset.

The non-weighted model with the intervention effect performed significantly worse, with an elpd_loo of -4743.58 and an elpd_diff of 211.34. This substantial difference underscores the importance of weighting observations by the number of games played, as it significantly enhances the model’s predictive accuracy. The non-weighted null model had the lowest elpd_loo at -4745.00 and the highest elpd_diff at 212.76, indicating that it was the least effective model in predictive performance.

Overall, the weighted model with the intervention effect emerged as the most robust and reliable model, demonstrating the value of both the intervention effect and the weighting scheme in improving predictive performance.

Figure 6 presents the scatter plot of predicted versus actual ΔOPS values generated using the weighted model with the intervention effect. The plot demonstrates a clear positive correlation between the predicted and actual values, with a coefficient of determination *R*^2^ of 0.52. This *R*^2^ value indicates that the model explains 52% of the variance in the true ΔOPS.

**Figure 6:**
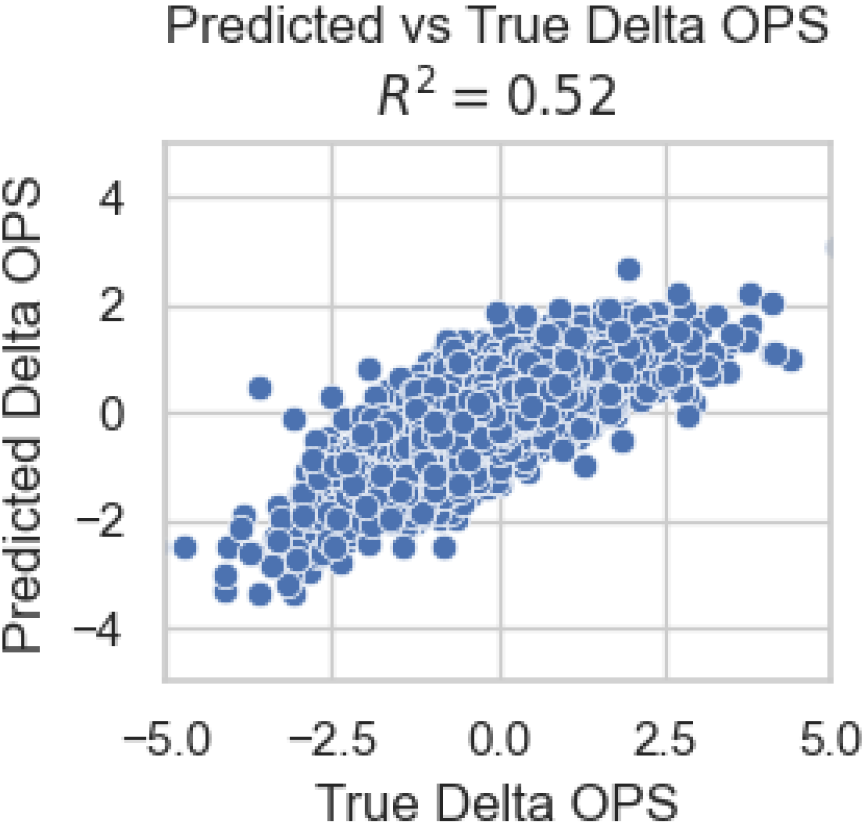
A scatter plot of the actual versus predicted change in OPS from the weighted model. The model had an R^2^=0.52.

### Previous Year’s OPS (β)

Figure 7A shows the posterior distribution for the previous year’s OPS coefficient (β). The distribution is centered around -0.758, with a 94% Highest Density Interval (HDI) ranging from approximately -0.783 to -0.735. The red vertical line represents the posterior mean of β. The negative value of β indicates that higher previous OPS values are associated with a decrease in ΔOPS, reflecting a diminishing returns effect where it is harder to achieve further improvement from an already high OPS baseline.

**Figure 7:**
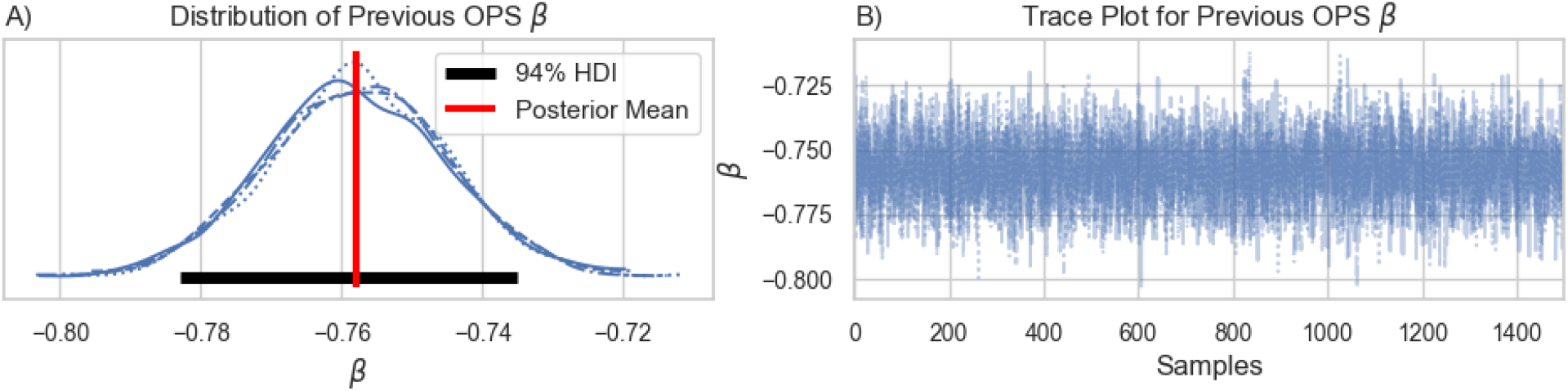
Weighted Model *β* coefficient results. A) Posterior distribution showing the mean (red) of -0.758 and the 94% Highest Density Interval (black). B) The posterior trace plots for the *β*_intervention_. All four chains are plotted in both plots, showing the high convergence of the model.

Figure 7B displays the trace plot for β, showing the sampling progression over iterations. The trace plot exhibits good mixing and stability, with no apparent trends or divergences, indicating that the MCMC chains have converged well. The 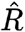 statistic for β was approximately 1, confirming the model’s convergence.

### Intervention Effect (*β*_intervention_)

Figure 8A presents the posterior distribution for the intervention effect coefficient (*β*_intervention_). The distribution is centered around 0.749, with a 94% HDI ranging from approximately 0.34 to 1.14. The red vertical line represents the posterior mean of *β*_intervention_. The positive value of suggests that the intervention has a significant and beneficial impact on ΔOPS, improving the team’s performance.

**Figure 8:**
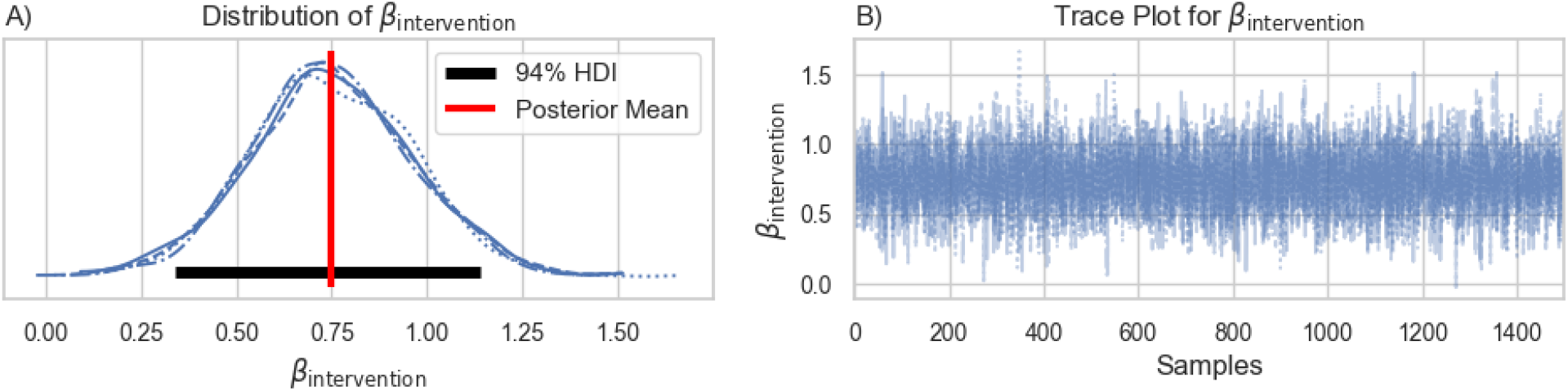
Weighted Model *β*_intervention_ coefficient results. A) Posterior distribution showing the mean (red) of 0.749 and the 94% Highest Density Interval (black). B) The posterior trace plots for the *β*_intervention_. All four chains are plotted in both plots, showing the high convergence of the model.

Figure 8B shows the trace plot for *β*_intervention_. Like the trace plot for β, this plot demonstrates good mixing and stability, indicating that the MCMC chains for *β*_intervention_ have converged well. The 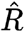 statistic for *β*_intervention_ was approximately 1, confirming the reliability of the posterior estimates. In addition, we computed the Bayes Factor for the *β*_intervention_ coefficient. The Bayes Factor (Kass & Raftery, 1995) was 72.30, indicating strong support for the intervention effect.

### *α*, Division, Year-Division, and Team Effects

The posterior distribution of the intercept, α, has a mean of -0.006 with a standard deviation of 0.823, and its 94% Highest Density Interval (HDI) ranges from -1.566 to 1.497. This indicates that, after accounting for other variables in the model, the baseline change in OPS is centered around zero, with considerable uncertainty.

The division effect has a mean of -0.103 and a standard deviation of 0.182, with a 94% HDI ranging from -0.432 to 0.260. This suggests that the division itself has a small and insignificant effect on the change in OPS, as the interval includes zero.

The year-division interaction effects show notable variability for specific years and divisions. For example, in 2016, the NCAA division had a significant negative effect with a mean of -2.315 and a 94% HDI from -3.306 to -1.317, indicating that this year and division substantially impacted the change in OPS. In contrast, the NAIA division in the same year had a much smaller and less significant effect, with a mean of -0.132 and a 94% HDI from -0.791 to 0.520.

Similarly, in 2020, the NCAA division had a substantial positive effect with a mean of 2.837 and a 94% HDI from 1.666 to 3.993, highlighting that this particular year and division experienced a significant increase in OPS changes. Other years, such as 2021 for the NCAA division, also showed a notable negative effect with a mean of -0.800 and a 94% HDI from -1.491 to -0.132.

These year-division interaction effects demonstrate that certain years and divisions have unique impacts on the change in OPS, which may reflect specific events or conditions affecting performance during those times. The model effectively captures these nuanced effects, providing a comprehensive understanding of how different factors influence team performance over time.

The team effects analysis revealed significant performance variability across different collegiate softball teams. After accounting for other variables in the model, these effects measure how much each team’s performance deviates from the average team performance. Teams with higher positive values of performance will have a higher ΔOPS compared to those with lower values. Oklahoma (NCAA) (4.06), Morris (S.C.) (NAIA) (3.92), Oklahoma City (NAIA) (2.92), James Madison (NCAA) (2.57), and UCLA (NCAA) (2.41) had the highest team effects, indicating significantly better performance than the average team. These high values suggest that these teams have consistently strong performances, possibly due to better training facilities, higher quality coaching, recruitment of top talent, and effective team strategies.

NAIA Team #1 had a team effect of 1.25, showing a significantly above-average performance. This indicates that NAIA Team #1 performed better than the average team, highlighting their year-to-year performance relative to peers. Positive team effects for NAIA Team #1 may be due to effective coaching, strong recruitment, and/or robust support systems for athletes.

NCAA Team #1 had a team effect of 0.54, indicating a better-than-average performance. This places NCAA Team #1 among the teams with positive deviations from the average, demonstrating solid performance. With a team effect of 0.32, NCAA Team #2’s performance was above average, though not as high as the other intervention teams. This suggests that NCAA Team #2 also performed better than average, but to a lesser extent.

The team effects analysis highlights significant performance disparities among collegiate softball teams. Teams like Oklahoma, Morris (S.C.), Oklahoma City, James Madison, and UCLA exhibited outstanding performance. The intervention teams showed better-than-average performances, with NAIA Team #1 leading. These results provide valuable insights into the relative performance of different teams and suggest that resource availability and organizational effectiveness are key factors influencing team success.

### Permutation Results

Figure 9 illustrates the permutation test results conducted to assess the significance of the intervention effect on the change in OPS (ΔOPS). The histogram in the figure shows the distribution of the permuted intervention effects generated by randomly shuffling the intervention labels and recalculating the effect multiple times.

**Figure 9:**
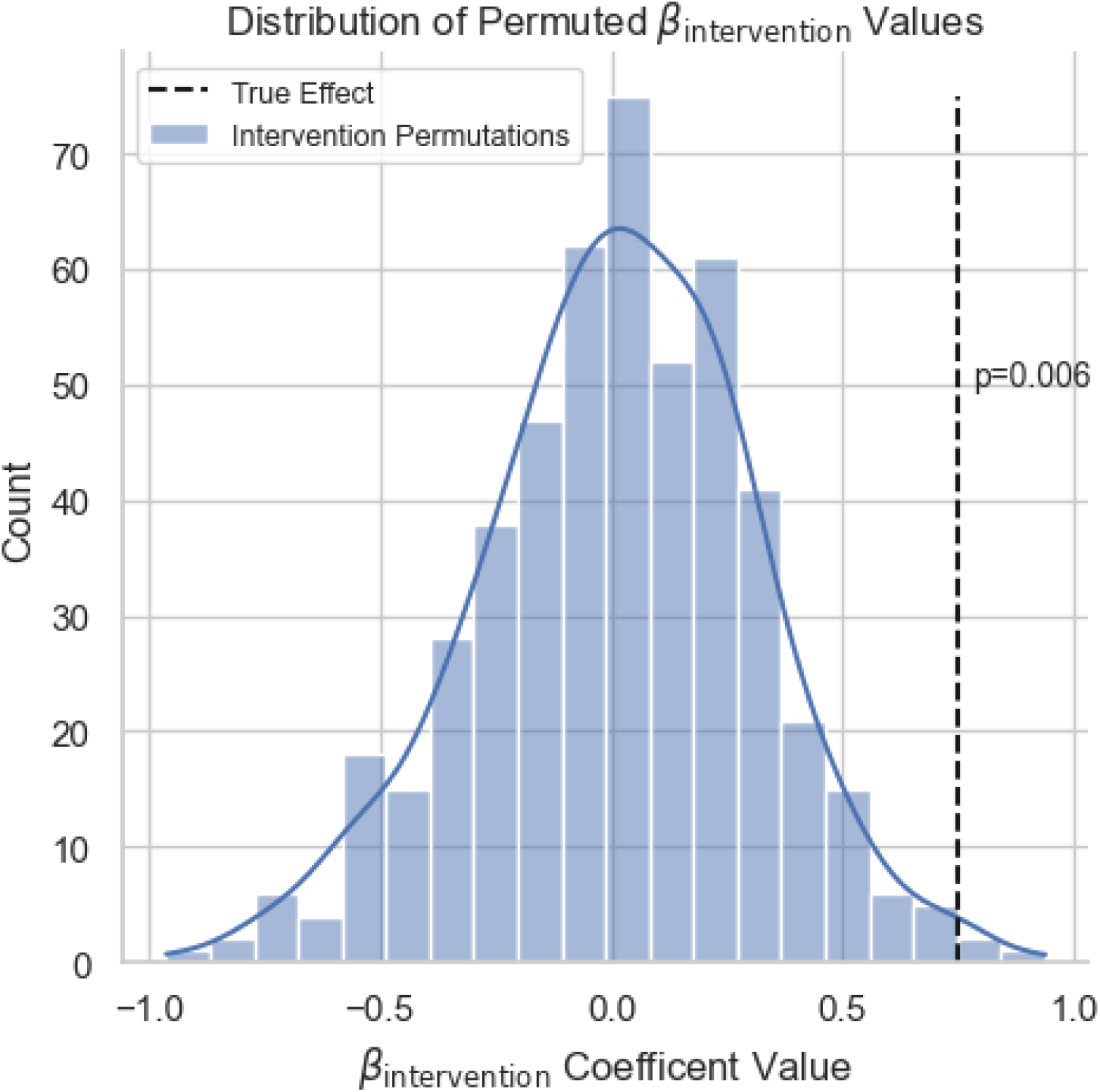
A histogram of the results of the permutation test. 500 permutations were run to test the randomized magnitude of the *β*_intervention_ coefficient. The real *β*_intervention_ coefficient is marked as the black dotted line with a p-value of 0.006.

The observed intervention effect, indicated by the red vertical line, is 0.749. This value is significantly higher than most permuted intervention effects, demonstrating a clear positive shift. Most permuted effects are centered around zero, suggesting that the observed positive effect is not due to random chance.

The p-value, calculated as the proportion of permuted effects greater than or equal to the observed effect, was found to be 0.006, further confirming the intervention’s statistical significance. This result provides strong evidence that the intervention had a meaningful and beneficial impact on team performance, as measured by the increase in OPS.

Overall, the permutation test validates the effectiveness of the intervention, highlighting a significant improvement in team performance that is unlikely to be attributed to random variation. The clear distinction between the observed effect and the distribution of permuted effects underscores the robustness of the intervention’s positive impact.

## Discussion

The findings from this study highlight the efficacy of targeted perceptual-cognitive training in improving the on-field performance of collegiate softball teams. The consistent improvement in Decision AUC and Response Time underscores customized training interventions’ significant role in enhancing specific batting skills. This is further supported by the permutation test results, which indicate a statistically significant positive effect of the intervention on team performance, as measured by OPS.

The substantial differences in training intensity among the teams, illustrated in the histograms of training pitches, suggest that the duration and frequency of training sessions are crucial factors in realizing these performance gains. Notably, NAIA Team #1’s more extensive training regimen correlates with more pronounced improvements in perceptual-cognitive assessments and real-world performance metrics. These results align with existing literature, emphasizing the importance of repeated, deliberate practice in mastering complex sports skills (Ericsson et al., 1993).

The uniqueness of this analysis lies in its longitudinal approach and the incorporation of assessment and training data alongside real-world performance metrics. By evaluating the effects of perceptual-cognitive training over multiple seasons and across different collegiate levels (NCAA and NAIA), this study provides robust evidence of the transferability of cognitive training to on-field performance. This approach contrasts with much of the existing literature, which often focuses on short-term interventions or laboratory-based outcomes, thereby limiting the ecological validity of their findings (Farrow & Abernethy, 2002). The present study’s use of a real-world, multiyear dataset enhances the external validity of the findings, suggesting that targeted cognitive training can sustain performance in competitive sports settings.

Moreover, the inclusion of weighted models that account for the variability in games played further strengthens the reliability of the findings. By giving greater influence to teams with more extensive in-season participation, the analysis more accurately reflects the conditions under which these interventions were implemented. This methodological consideration addresses a common limitation in sports performance research, where unequal participation rates can skew results (Ericsson et al., 2018). Future studies could build on this approach by exploring the differential impacts of cognitive training across various positions or roles within a team and investigating the optimal frequency and duration of training sessions to maximize on-field performance improvements.

In addition, this study adds weight to the growing body of literature indicating cognitive-perceptual training in sports needs to be ecologically valid. Previous studies have shown that cognitive training can enhance decision-making speed and accuracy in athletes (Mann et al., 2007). Still, few have provided direct evidence of how these improvements translate into enhanced performance metrics like OPS (Fransen, 2024). The observed improvement in OPS, particularly among teams that engaged more intensively with the training, suggests that cognitive training enhances perceptual skills and leads to measurable gains in competitive outcomes. This finding aligns with the theory of cognitive transfer, which posits that skills developed in one context (e.g., a training module) can transfer to another (e.g., a game situation) if the tasks share underlying cognitive processes (Barnett & Ceci, 2002). More importantly, the cognitive training used here – the uHIT Platform – is ecologically similar to the decision process in the field.

Future research could expand on these findings by examining the long-term effects of cognitive training after the intervention’s cessation. Such studies could assess whether the benefits observed in this study persist in subsequent seasons without continued training or if maintenance sessions are necessary to sustain the gains. Additionally, investigating the effects of cognitive training across different sports or among different levels of competition could provide further insights into the generalizability of these interventions.

Overall, this study underscores the potential of cognitive training as a critical component of athletic development programs, particularly in sports where perceptual and decision-making skills are paramount. The results suggest that integrating such training into regular practice routines could offer teams a competitive edge, particularly when combined with traditional physical training methods.

## Acknowledgments

We’d like to thank the NCAA and NAIA teams for their support and use of the uHIT Platform. Research was sponsored by the Army Research Office and was accomplished under Grant Number W911NF–23–1–0271. The views and conclusions contained in this document are those of the authors and should not be interpreted as representing the official policies, either expressed or implied, of the Army Research Office or the U.S. Government. The U.S. Government is authorized to reproduce and distribute reprints for Government purposes notwithstanding any copyright notation herein.

